# Evolution of simple multicellularity increases environmental complexity

**DOI:** 10.1101/067991

**Authors:** María Rebolleda-Gómez, William C. Ratcliff, Jonathon Fankhauser, Michael Travisano

## Abstract

Multicellularity—the integration of previously autonomous cells into a new, more complex organism—is one of the major transitions in evolution. Multicellularity changed evolutionary possibilities and facilitated the evolution of increased complexity. Transitions to multicellularity are associated with rapid diversification and increased ecological opportunity but the potential mechanisms are not well understood. In this paper we explore the ecological mechanisms of multicellular diversification during experimental evolution of the brewer’s yeast, *Saccharomyces cerevisiae*. The evolution from single cells into multicellular clusters modifies the structure of the environment, changing the fluid dynamics and creating novel ecological opportunities. This study demonstrates that even in simple conditions, incipient multicellularity readily changes the environment, facilitating the origin and maintenance of diversity.

## Introduction

A world without multicellular life is a world without trees, coral reefs, nests and cities. A unicellular world lacks most of the architectural complexity we are familiar with. There is no doubt that multicellular life has deeply transformed the world around us. The integration of previously autonomous cells into a multicellular organism involves major changes of organization. This architectural complexity is often associated with the evolution of novel ecological opportunities, but the ecological consequences of this spatial reorganization have been largely overlooked. Even simple multicellularity involves changes in the arrangement of cells, imposing new spatial structure. This spatial heterogenetity has been deemed important for the evolution of cellular differentiation [1] but its consequences for the origins and maintenance of diversity remain largely unexplored.

Multicellularity, has evolved multiple times with varied consequences for subsequent diversification and complexity [2, 3]. However, even relatively recent origins of multicellularity occurred more than 200 MYA [4]. Therefore, direct observations of its ecological consequences are necessarily limited in natural populations.

*Saccharomyces cerevisiae* rapidly evolve multicellular phenotypes in the laboratory under settling selection, providing an experimental model for direct investigation of the effects of this reorganization for the origin and maintenance of diversity [5, 6]. Ten replicate populations originated from a single diploid isogenic unicellular strain. Every day a subsample of each culture was taken and centrifuged at very low speed, thereby increasing the representation of large individuals at the bottom of the tube. Following centrifugation, the bottom 100*μ*l were transferred to fresh media and allowed to regrow over 24 hours. After 60 days of selection, all populations evolved multicellular phenotypes [5]. In this system (“snowflake yeast”), post-division adhesion of mother and daughter cells limits the potential for among-cell conflict and confers multicellular heritability, facilitating multicellular adaptation [5, 7]. The simplicity of this system allows us to carefully parse the ecological consequences of multicellularity and their effects on the observed diversity.

Simple evolutionary models, assuming a smooth and unchanging adaptive landscape [8], predict convergence with respect to size, because size is tightly correlated with fitness during settling selection [5, 6]. In contrast, high levels of heritable phenotypic variation for snowflake cluster size evolved both within and among replicate populations. Most of the diversity was observed among replicate populations. Nevertheless, after 60 days of selection, nine of the ten populations harbor two or more genotypes that differ in average cluster size, suggesting that the evolution of multicellularity also facilitated morphological diversification within populations. This paper investigates the ecological dynamics of incipient multicellularity in one of these populations and its consequences for rapid diversification [6].

## Results

### Phenotypic diversification

Focusing on the above mentioned replicate population (C1) after one, four and eight weeks of settling selection, we determined the size distributions of ten randomly chosen single colony isolates at each time point. Diversity in cluster size could in principle arise from one or more evolutionary processes: neutral divergence, selective sweeps, and adaptive diversification. These processes differ in the tempo of evolution, and can be partially distinguished by determining the rate of morphological diversification [9, 10]. In this population, multicellularity evolved within one week of selection, but the clusters were relatively small, initially rare (1 in 10 isolates; Fig.1A), and had a relatively minor increase in settling rate [5]. After four weeks of selection, seven of ten isolates were multicellular and at least two different size classes had evolved (Fig.1A). Three of the multicellular isolates had decreased size, but the other four were four-fold larger in size (Fig.1A, Fig. S1A). Following 60 days of selection in total, two clearly distinct life-histories had evolved with evidence of incipient diversification within each life-history (Fig.1, Fig. S1B)(one-way ANOVA, effect of strain F(9, 20) = 103.26, *p* – value < 10^−^14). This pattern of rapid diversification and persistent diversity in fitness-related traits is most consistent with increased ecological opportunity leading to adaptive diversity and not with neutral divergence or selective sweeps.

**Figure 1:**
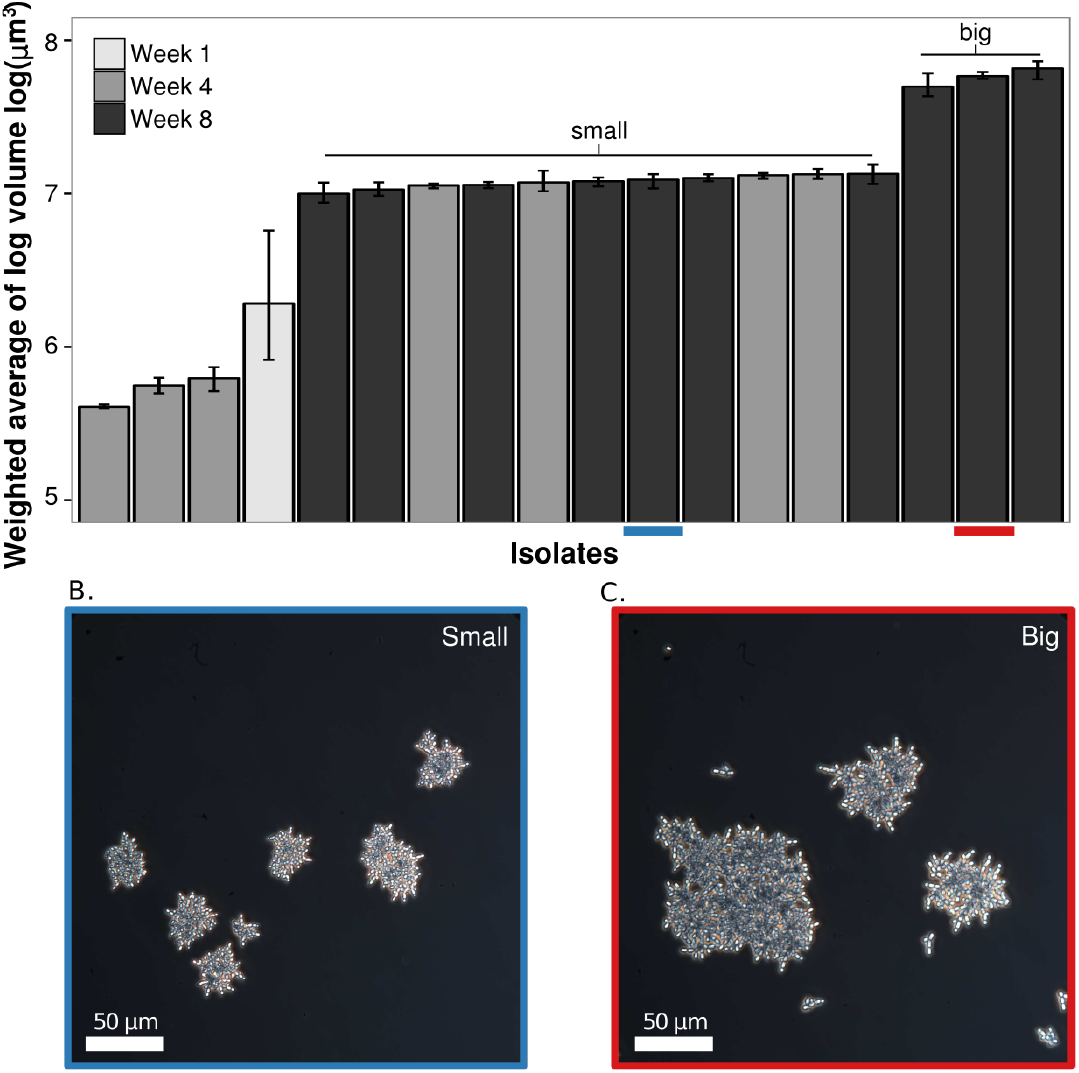
Diversity on size distributions of isolates from one of the replicate populations. (A) Volume (weighted by total biomass) of different multicellular isolates throughout the duration of the selection experiment. Each value is the average of three replicates, each composed of six different fields of view and at least 70 individuals. Big and Small groups can be recognized statistically based on a Tuckey HSD test with significance level of 0.01. Error bars are 95% confidence intervals. (B, C) Pictures showing typical adult and propagule size of the two main strategies described:Big (red) and small (blue).

To confirm adaptive divergence as the underlying process, we investigated the ecological basis for maintaining phenotypic size variation within populations. The two primary determinants of competitive fitness during the experimental evolution were growth and settling rates. We measured both of these components on two representative isolates from population C1 after 8 weeks of selection. Differences in growth rates were observed among the two isolates (Table S1, Fig. S2).

We determined settling rates that incorporated maximal and minimal cluster sizes of an isolate, reflecting cluster size immediately prior to reproduction and the minimal size of the resulting offspring, respectively. Settling rate differences were measured by changes in optical density during settling after 24hrs of growth (Fig.3D). Reduction in absorbance follows a sigmoidal curve that can be approximated by a logistic equation with *K_OD_* as the inflection point (the time at which there is a half reduction in optical density) and S as the settling rate or steepness of the curve (Fig S2C). Isolates with large cluster sizes (“Big”) settle rapidly (larger S value; Fig.3A, Fig. S2C, Fig. S3A). Multicellular isolates diversified into at least two life-histories: one encompassing clusters with a large adult size and fast settling rates that reproduce into many small propagules (Fig.1, Fig.3A). These isolates have a long juvenile phase and pay a cost in terms of their overall growth (Fig S2B, Video S1). The second life-history includes isolates with smaller adult sizes and a reduction of the juvenile phase with larger propagules (Fig.1, Fig. S1B, Fig. S2B, Video S1).

### Coexistence is stable

Coexistence between these two life-histories could be transient–the result of an incomplete selective sweep. To evaluate the stability of this relation we performed a series of competition experiments with one of the small and rapid growing genotypes (“Small”) against a “Big” genotype incorporating an inducible, effectively neutral GFP marker. The marker has no detectable fitness effects in this environment (two sided t-test, *t*(31.09) = 1.5292, *p* – *value* = 0.1363, mean diff= 0.678, 95% CI [−0.02029751, 0.14200565]). We performed these competitions under different intensities of settling selection. As expected, under very intense selection (3 min of settling), the big and fast settling strain rapidly increases in frequency, almost displacing the smaller strain after only three transfers. In contrast, relaxed selection (25min of settling) favors the small but rapid growing strain. Coexistence, nevertheless is possible at intermediate selection (7 minutes of settling) and the frequencies remain stable over the course of this assay (Fig. 2A). Stable frequencies at intermediate selection and ecological differences between these strains are inconsistent with the hypothesis of an incomplete selective sweep.

**Figure 2:**
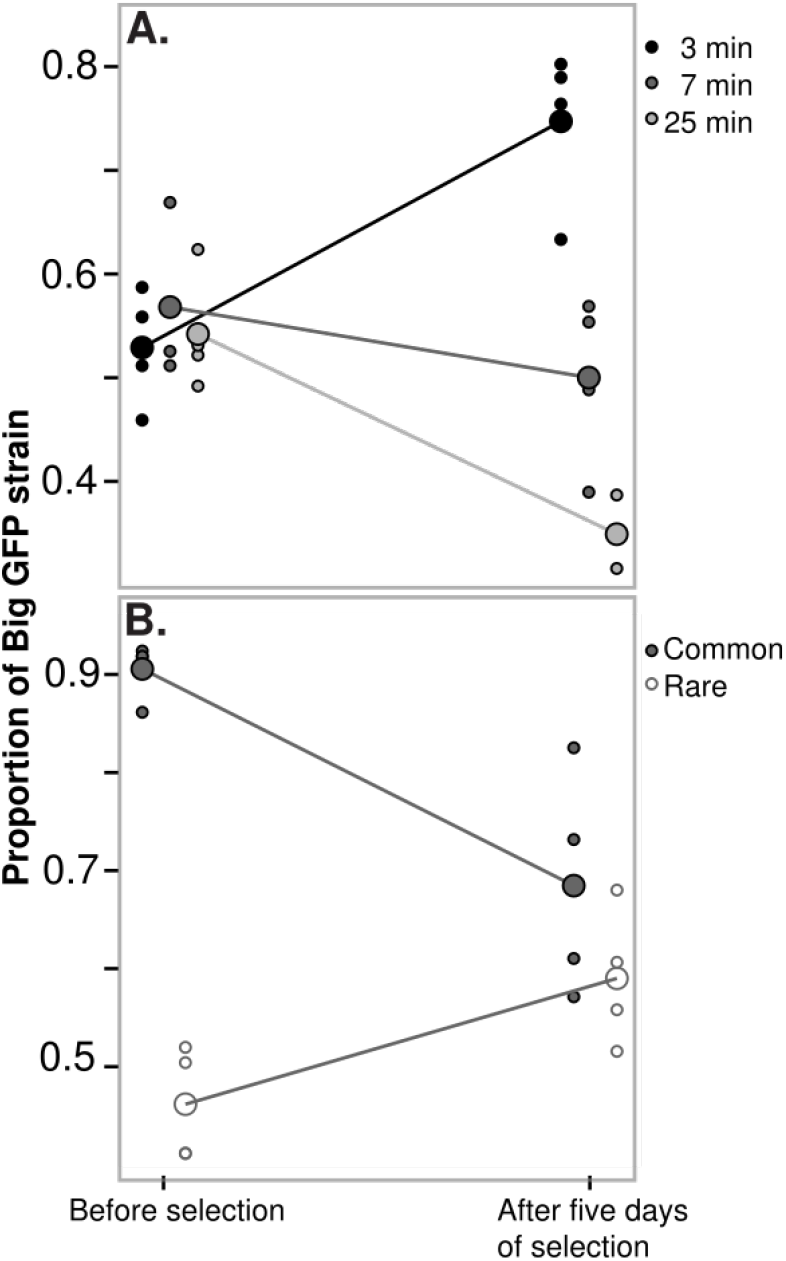
Competition assays. Both plots display the frequency of the “Big” genotype with a GFP marker before and after 5 days of settling selection. (A) Intense selection (3 min of settling) favors the fastest settling strain (“Big”), relaxed selection (25 min of settling) favors the fastest grower (“Small”) and coexistence is maintained at intermediate selection (7 min of settling).)(B) At intermediate selection both genotypes exhibit negative frequency-dependent fitness and therefore can invade from rare. Small points show the data of each of the four replicate competitions in each treatment and the large point and lines represent mean values.

To completely rule out this possibility we performed a second competition experiment under settling selection of intermediate strength (7 min). In this case, we started with each of the strains at a low frequency and asked if they could invade. Both strains increase in frequency when rare, reaching a stable 6:4 ratio after five days of competition (Fig.2B). Together, these results suggest that coexistence between these two strains is due to negative frequency dependent dynamics resulting from a trade-off between growth and settling, with intense selection favoring the larger strain and relaxed settling selection favoring the faster growing but smaller strain. We used computer simulations to investigate the potential for this trade-off to explain the patterns observed.

### Trade-offs alone fail to explain coexistence

We investigated whether the observed growth and settling rate differences were sufficient to maintain coexistence. In this environment, larger clusters have an advantage because they settle faster. However, there is a trade-off and larger genotypes pay a cost in terms of growth rate [5, 6](Fig.3A, Fig.S2). Using settling and growth data we simulated competition between strains. In YPD medium (the one used throughout the experiment) *S.cerevisiae* displays diauxic growth, changing from exponential to nearly linear growth (Fig S2A). Using piece-wise regression models we estimated the slopes and breakpoints for each strain (table S1). Competition was included in our simulations by defining the points of growth-rate change for each strain in terms of the overall culture density (Fig.3B). In the absence of settling selection, growth rate differences alone allow the fast growing strains to outcompete the large, rapidly settling isolates (Fig.3C). Incorporating settling selection (by adding the proportion of individuals that would settle in a given time)(Fig.3B,D) in the model increases the frequency of the big strain initially, but it is not sufficient to promote coexistence (Fig.3C).

**Figure 3:**
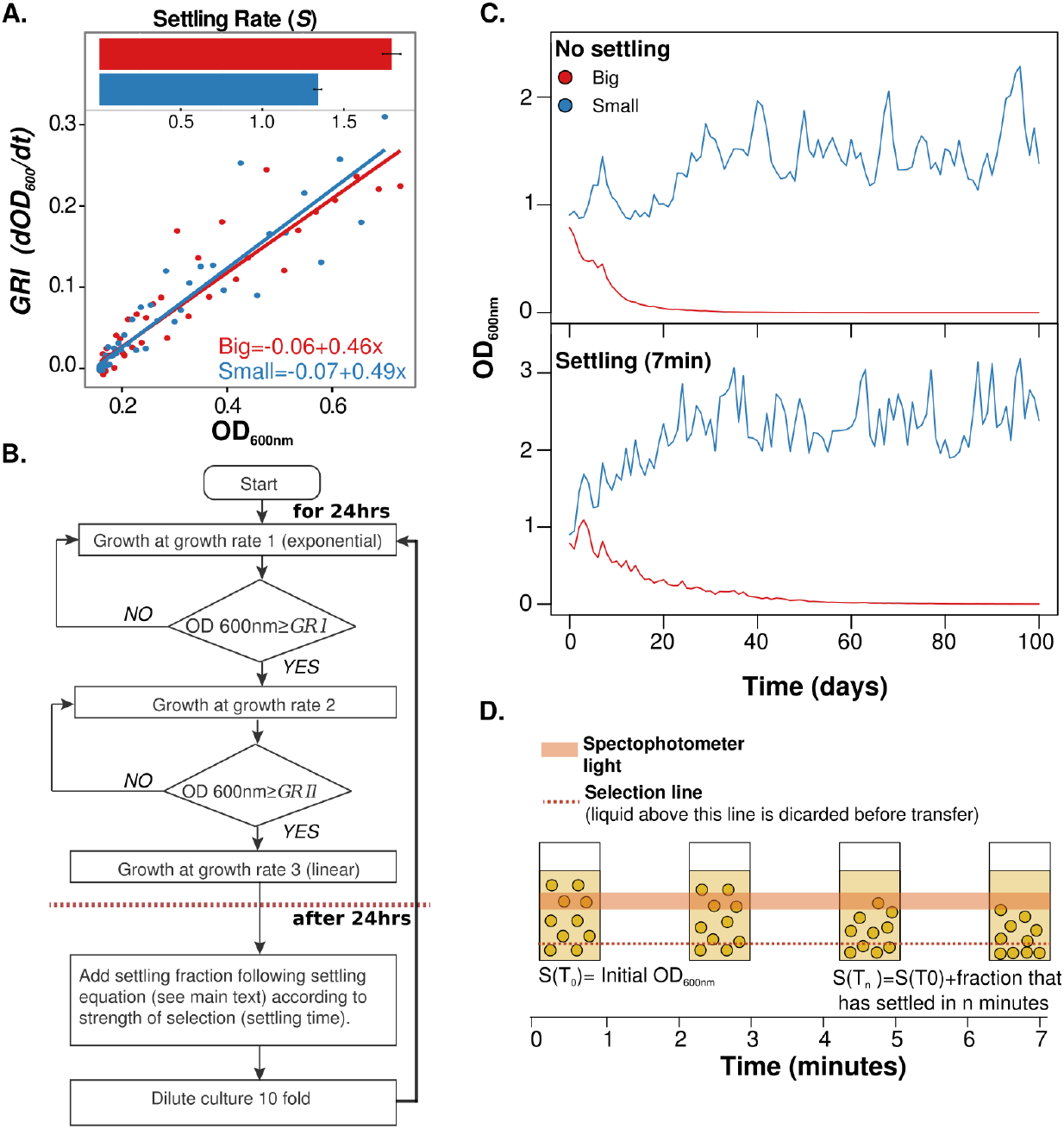
Simulations of competition between the big and small strains.(A) There is a trade-off between growth and settling. Differences in slopes of growth rates against 0D at 600nm during the exponential phase are small but enough for the small fast-growing strain to displace the big strain in our simulations of competition. Each point was calculated from six replicate growth curves. Settling data is the average estimate from six replicate settling assays and the error bars represent one standard deviation. (B) Algorithm for simulations. Using a segmented regression, changes in growth rates are estimated. The minimal OD at which any of the strains changes their growth is set as the point for both to change their growth rates (*GRI* and *GRII*), settling is added as after 24hrs of growth using a logistic equation (D; see methods). (C) A representative round of simulations of competition over 100 days without settling (upper panel) and with 7 min (intermediate selection) of settling (lower panel). Our simulations fail to recover coexistence between these two strains despite large differences in settling rates (A-small panel).

This simulations show that even very small costs in terms of growth rate are able to lead to competitive displacement over the long term (Fig.3). Given the small differences we were unable to find statistical differences in growth rates calculated from OD at 600nm data. In previous studies, however, larger differences in growth have been observed by measuring changes in total biomass [6]. Our simulations were built with these conservative estimates of growth differences,and nevertheless, the Big strain is not able to recover even in the presence of settling selection.

Our *in vivo* competition assays suggest that the above simulations do not include all the ecological complexity involved in the interactions between different members in this environment. These assays provide evidence for the importance of frequency-dependent mechanisms in the coexistence of these two strains. The role of multicellularity in promoting diversification in this system cannot be understood in terms of a tradeoff between growth and settling; this novel organization alters the ecological dynamics in a non-additive manner. What are the mechanisms underlying this frequency-dependent interaction?

### Increased environmental complexity allows for coexistence

We hypothesize that at first, when big individuals are rare, they settle faster than smaller clusters. However, once there is a high frequency of clusters in the media, the effective viscosity of the medium also increases; sinking of an individual depends on its displacing not only the liquid but also other suspended individuals slowing down their descent. This phenomenon of hindered settling is well known for dense sediments and its effects are more intense towards the bottom—where particles concentrate [11]. In these same conditions of high density, smaller individuals are more likely to percolate, increasing the chances of these smaller clusters being transferred to the next tube.

To test these hypotheses about the effects of each strain’s frequency on the settling dynamics, we mixed the fast growing (“Small”) and fast settling (“Big”) strains in different proportions. We then measured the median cluster size of the culture before selection as well as the median cluster size of the bottom fraction after seven minutes of settling. When the mixture is initially composed of mainly small clusters (small median size) only the biggest clusters reach the bottom in the time provided. As a result, there is an overall increase in size after settling. By contrast, when the culture has a larger proportion of bigger clusters, a disproportionate number of small individuals survive settling selection, resulting in a net decrease in size (Fig.4A).

**Figure 4:**
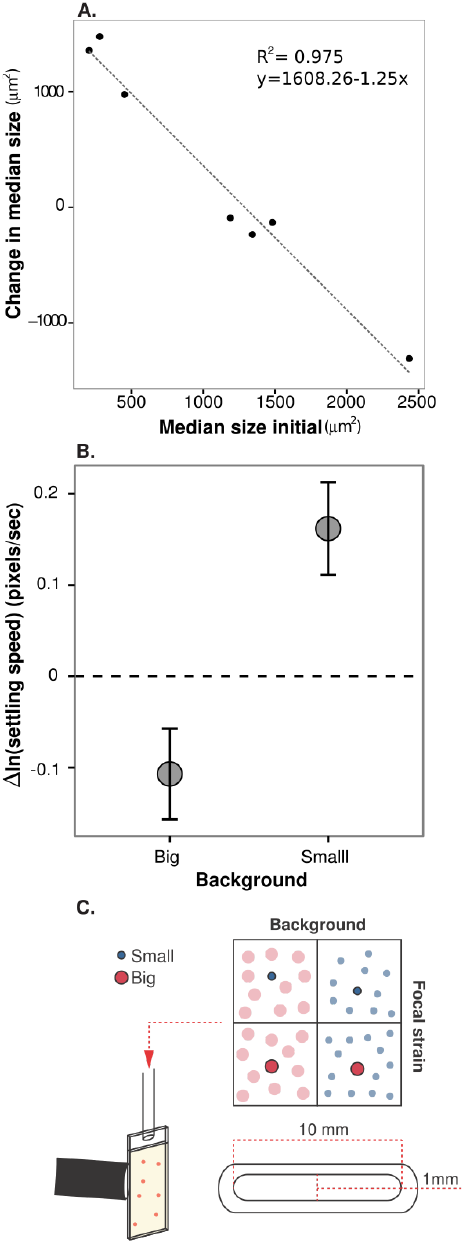
Increased environmental complexity and settling interactions result in frequency dependent dynamics. (A) Change in median size after settling for a range mixes with different proportions of “Big” and “Small” and thus different initial median sizes. From each mix we sampled six fields of view and at least 58 clusters for the initial time point and at least 127 clusters after settling. (B) Difference in log transformed average settling speeds between the Small and Big strains in both backgrounds (Big-Small). Differences are significant and in different directions: on average, clusters of the Small strain settle faster in the Big background and Big clusters in the Small background. The error bars represent the 95% confidence intervals. We performed 6 replicate settling assays with different cultures of the same strains, following clusters settling for approximately 2min, with at least 82 clusters tracked per replicate, per treatment. (C) Experimental setup to measure settling speed in different strain backgrounds. A focal strain stained in red with safranin was mixed in a 1:10 ratio (in volume) with a dense overnight culture of either the same strain or the other one. Images were recorded every 0.1s as the culture settle within a thin glass chamber.

To directly measure the effects of high density (*i.e.* increased effective viscosity, and drafting or reduced drag for smaller particles) on settling speed, we recorded the speed of clusters of each strain (“Small” or “Big”) as they settled with a majority of clusters of the same or the other strain (Fig.4C). Drafting and increased effective viscosity act in different directions and their relative importance depends on the density and the properties of the particles [11]. To control for the effects of density, we performed our experiments with dense cultures after 24hrs of growth (the same density as when experiencing settling selection before each transfer). In high density backgrounds (“Big” and “Small”) settling is hindered slowing down each cluster with respect the same focal strain density in the media (YPD) background (i.e. settling in a low density treatment; data not shown). We found an effect of background and background focal-strain interaction (Table S2). Consistent with drafting, both strains settle faster when they are in a big strain background (planned contrast “Big” as focal-strain, *z* = 4.195, *p — value* < 10^−^4; “Small” as focal strain, *z* = 19.595; *p – value* < 10^−^4B). However, there is more variation in the big strain background as would be expected from increased effective viscosity and a wider distribution of particle sizes (Fig. S1B; Fig. S3).

Due to hindered settling, in the big background we would expect that the largest clusters will still be likely to settle faster, but small and dense clusters are more likely to percolate through spaces between big clusters slowed down by the increase in effective viscosity. This effect will tend to favor the small strain: given the same cluster size, individuals of the smaller strain tend to be more dense and circular (Fig.5B). Consistent with these hypotheses and our previous observations, the “Big” genotype settles slower (relative to the smaller strain) in its own background (planned contrast “Big” background, *z* = –5.282, *p* – *value* < 10^−^4). Instead, in the small background, larger clusters are able to displace their smaller competitor (planned contrast “Small” background, *z* = 7.843, *p* – *value* < 10^−^4; Fig.4B), explaining their advantage when rare.

**Figure 5:**
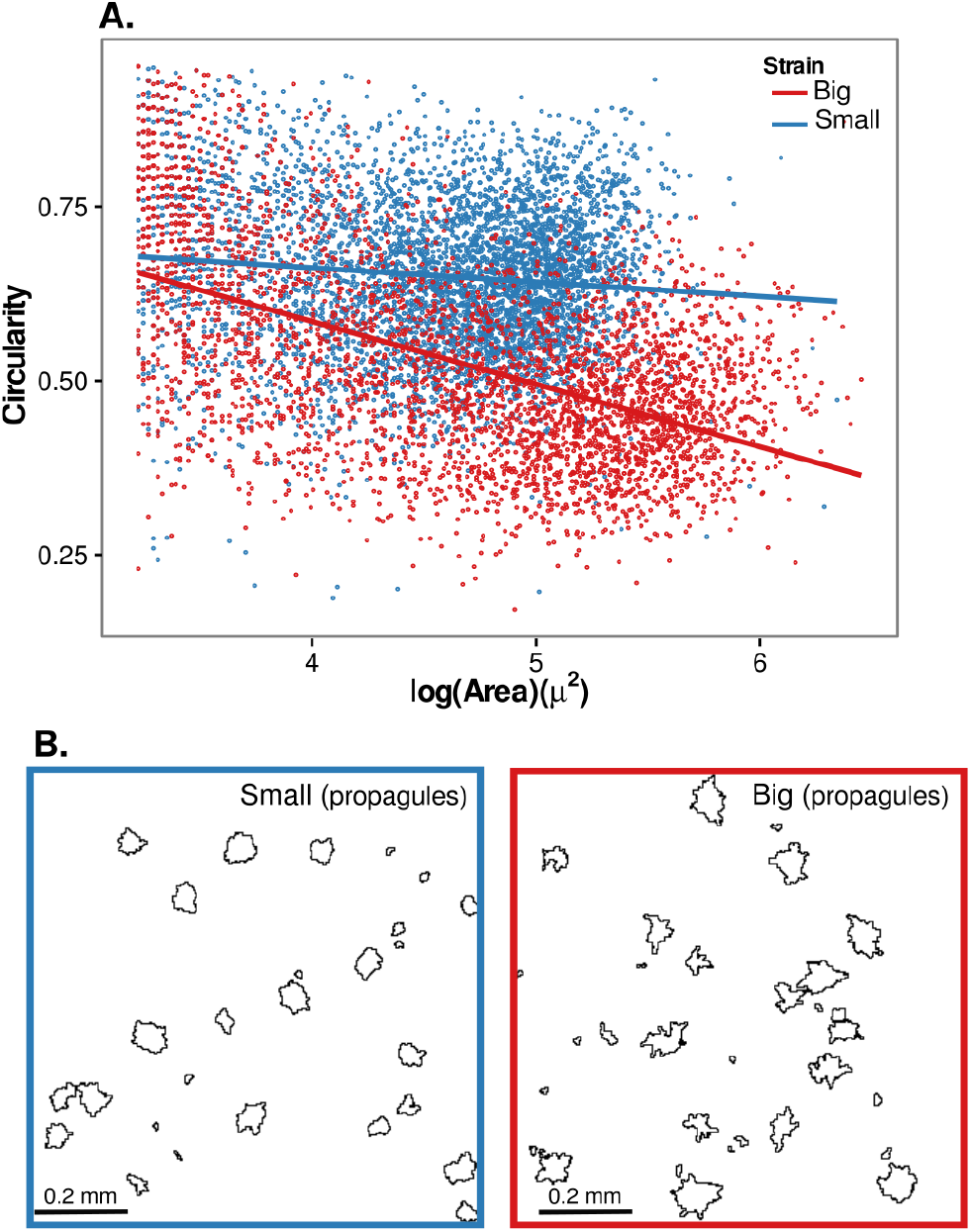
Clusters of the same size tend to be more circular and solid in the Small genotype. (A) Relation between size and circularity for each strain. (B) Outlines of propagules of similar sizes. Small clusters of the small strain tend to be better packed and more circular.

## Discussion and conclusions

Evolutionary innovations expand the phenotypic space, allowing for the exploitation of resources in ways that were previously not possible [12–14]. The evolution of multicellular yeast phenotypes increased the diversity of phenotypic possibilities of *S. cerevisiae* under the laboratory conditions. The multicellular isolates show variation in traits involving multiple cells like cluster size, number of propagules, adult sizes and the size of propagules. These qualitatively distinct traits—changing the space of ecological and evolutionary possibilities—do not always lead to diversification, and the role of innovations is highly contingent on ecological processes like availability of ecological opportunity [14–16] and competitive dynamics [10,17,18]. However, Key innovations—like the evolution of multicellularity in this experimental system—can have a greater impact by changing their own (and their competitors’) environment, transforming, in this way, selection pressures and creating novel ecological opportunity. It has been proposed that innovations involved in this processes of “ecosystem engineering” might have a greater impact on the origins and maintenance of diversity [10,15,19].

Coexistence of the two groups identified by size in this paper, arises largely due to ecosystem engineering. Growth modifies the environmental conditions making them more favorable for the least common type, enabling coexistence in a frequency-dependent manner. Ecological theory has shown that coexistence in temporally varying environments with a single limiting resource is possible if species differ in their resource use at different concentrations of the resources [20, 21]. Consistent with these models, all the isolates evaluated in this work displayed a diauxic growth pattern (reflecting a non-linear dependence on resources) and different isolates evolved different non-linear dynamics (Table S1, Fig. S2A). However, our simulations show that these differences in growth were not sufficient to promote stable coexistence. In this system, additional non-linear effects are due to the complex dynamics of settling, and our simulations suggest that these added effects can be important for coexistence of additional types varying in adult size (Supplementary text, Fig. S3). These observations suggest that complex interactions between different mechanisms of coexistence might be at play, allowing for increased variation in different components of these multicellular life-histories (Supplementary text, Fig. S3), with environmental modification being an important driver of this diversity.

Niche construction theories propose that by modifying their environment, organisms change their own fitness values affecting subsequent evolution [22]. It is well known that microbes often modify their environment by secreting by-products of their metabolism [23]. Ecological specialists can then arise taken advantage of these by-products as main resources. In this system, evolutionary integration of multiple cells in multicellular clusters modifies the spatial structure of the environment, changing the settling dynamics and leading to increased environmental complexity. These new dynamics affect the fitness values of individual clusters in a negative frequency dependent manner creating the opportunity for coexistence and diversification. Existing theoretical frameworks of niche construction [22] and extended phenotypes [24] most commonly refer to cases in which modifications of the environment are correlated with an increase in fitness for the organism. Evolution of environmental modification is favored because extended phenotypes are subject to positive selection [24]. In this case, environmental modifications are byproducts of size and population density (i.e. selection is not acting directly on hindered settling), and their fitness effects depend negatively on frequency. This system exemplifies the role of eco-evolutionary feedbacks promoting processes of rapid diversification [25] and provides evidence for the importance of “ecological byproducts” on evolutionary change.

It has been proposed that ecosystem engineering and environmental modifications were important in the Ediacaran-Cambrian radiation of metazoans, specifically biogenic mixing of nutrients by sponges and burrowing organisms [26]. This study provides evidence for the potential of multicellularity to modify the environment, increasing the ecological complexity and promoting diversification. This paper is an empirical demonstration of a theoretical framework, showing that multicellular organization allows for the construction of new ecological opportunity. This framework provides avenues for future research in the ecological and evolutionary consequences of multicellularity. The experiments in this study were performed in highly controlled environments, and the multicellular phenotypes described here are much simpler than most other multicellular forms. However, if in an environment as simple as this, with organisms as simple as the ones presented here, we can observe the evolution of such ecological complexity and diversification, one can well imagine the potential that the first multicellular forms had in a world that is immensely more complex than a test tube.

## Materials and methods

Unless specified otherwise, we grew all our strains in 10ml of Yeast Peptone Dextrose medium (YPD per liter: 10g yeast extract, 20g peptone, 20g dextrose) at 30°C and constant shaking at 250 rpm. All the strains used in this experiment were isolated from one of the ten replicate populations from [5]. Isolates from week one and four were isolated at random from frozen samples of one of the replicate populations (population C1) from weeks one, four and eight of the original experiment. At each of these time-points, we stroked the frozen sample on an YPD plate with 15% agar. We took a loop of culture and grow it for 24hrs. After a full day of growth with diluted the culture and plate-it to a total effective dilution of 1:100,000. After 48hrs of incubation on the plates, we selected at random 10 colonies (using a grid and a a sequence of ten random generated numbers) and streaked each of them on a new plate. To obtain single genotype colonies, we plated each of these strains in three consecutive plates, selecting a single colony and streaking it each time. Isolates from week eight were isolated previously, using the same method [6]. All isolates were then kept in 20% glycerol vials at −80°C. All strains were assigned a random number, the two strains chosen for the rest of the experiments were selected by sampling a random number from the id numbers of strains in each group (Big or Small).

### Size measurement and biomass calculations

To have comparable measurements of all isolates, all the strains were grown at the same time. After conditioning, strains were transferred again to 10ml of fresh YPD and grown for other 24hrs under the same conditions. We then placed 10*μ*l of a 1:10 dilution in a hemocytometer chamber. We photographed six fields of view using phase-contrast and the 40X objective on an Olympus IX-70 with a SPOT Flex 64 MP camera. The acquisition properties were set to maximize contrast between the yeast cells and the background and were kept constant throughout the experiment. Size of clusters was then determined using ImageJ [27]. We repeated this procedure with three replicate cultures per strain. Measures of unicellular isolates were not used in this study. In all of our microscopy work, to have a representative sample, each slide was divided in a grid of six sections and images were taken in one of each of these sections.

Instead of using raw measures of cluster size, for these analyses we calculated a weighted “biomass” measure. Distributions of cluster size can fail to predict the ecological dynamics and changes in how biomass survives settling selection. This is due to the fact that, for a given number of cells (or amount of biomass), the formation of larger clusters reduces the number of clusters that are counted. This is particularly problematic for the isolates with very large ‘adult’ multicellular clusters and many small ‘juveniles’: most of the biomass may reside in a small number of adults, but the statistics of the distribution will be dominated by traits of juveniles. We solve this problem by calculating the distribution of biomass within the population using a custom function in R [28](Software S1). To do this, we give each cluster a weight proportional to its biomass, calculating the mean fraction of the population’s biomass that falls within specific size range bins.

This weighted measure of volume, was log transformed to reduce the asymmetry in the distribution. We performed a one-way ANOVA with these log transformed values as the dependent value and strain as our factor. The ANOVA results were significant and post-hoc Tukey HSD tests were used to assign strains to distinct size classes.

Adult size comparisons were used to select representative strains for further analyses as described in [6]. The strains used were: the first multicellular strain identified in the experiment (C1W1.1), two strains from week four with different sizes (C1W4.1 and C1W4.3) and four strains from week eight representing the four size classes identified in [6] (C1W8.1, C1W8.2 (Big), C1W8.8, C1W8.9 (Small)).

### Determination of settling speed and growth assays

To compare the settling speeds of each isolate, we placed 1ml of a 24hr culture of each strain in a plastic cuvette and homogenized the culture before placing it in a spectrophotometer recording the absorbance at 600nm approximately every 15 seconds for 7 minutes. Each of the six replicates was independent (different yeast cultures were used for each replicate). Data was fitted to a sigmoidal curve with self-starting values using R [28, 29]. Given that each replicate had different starting densities, we fitted each replicate independently and then calculated the mean and standard deviation for the two main parameters of interest: the settling rate (*S*) and the inflection point (*K_OD_*).

We used a Tecan Infinite 200Pro plate reader with a 96 well BD Falcon flat transparent bottom plate to measure the optical density of our cultures at 600nm every 10 minutes over the course of 24 hours. Between measurements, the plate was shaken for seven minutes using the orbital mode with amplitude of 1mm we used linear shaking for the remaining three minutes. This combination allowed us to keep all clusters in suspension throughout the experiment. Temperature was kept constant at 29.9°C.

### Competition simulations

To simulate competition between the four isolates of interest, we first calculated their growth rates at each concentration (optical density at 600nm). And using a piecewise regression with the “segmented” package in R [28, 30] we estimated the breakpoints where changes of slope are significantly different from zero. We then estimated the slopes of the different fitted segments. These slopes where used as growth rates in our simulations. Growth rates of each strain depend on the overall culture density. Once the sum of all optical densities reaches each minimal breakpoint, all strains switch to the next growth rate (Fig.3B; Fig S2A). In the absence of selection, growth proceeds in this way for 24hrs and, after a day of growth, the density of all isolates is divided ten fold (simulating the dilution process between transfers with a fraction that allows for full recovery of density before the next transfer). Selection is incorporated after the 24hrs of growth and right before the dilution step. The fraction that settles to the bottom is calculated by incorporating the culture density after the desired settling time in the logistic equation fitted to the settling data. Settling at the bottom of the tube gives an increase in density over time at the same rate that density is reduced in the upper fraction of the tube. Given that our data on settling was obtained by measuring reduction in optical density in the upper fraction of the tube over time, to calculate the increase in density at the bottom of the tube we changed the sign of the slope (Fig.3D; Fig SC; Software S2).

### Competition assays

“Big” cells with inducible GFP fluorescence were created by replacing URA3 with *yeGFP* under the control of the *ScMET25* promoter [31] via the LiAc/SS-DNA/PEG method of yeast transformation [32]. Transformations were plates on solid YPD (15% agar) with 100mg/L of the antibiotic clonNAT. For each transformant, the insertion location of the transformation sequence was confirmed by PCR. The cost of the marker was evaluated through four days of competition without selection. In each of the three replicates, 50*μ*l of this “Big”-met25 GFP were inoculated with 50*μ*l of “Big” (no marker) in 10ml of YPD and grow for 24hrs before transferring 100*μ*l to fresh media. Frequencies were determined after 24hrs of growth and after four days of daily transfers. Change in frequency between the initial and four days was evaluated with a two-sided t-test.

The ”Big” strain with the GFP marker was mixed in a 50:50 ratio with the ”Small”. These strains were inoculated as a 1:100 dilution in 10ml of YPD. After 24hrs of growth at 30°C and 250rpm a sample was taken to determine the proportion of met25 GFP individuals before selection (T0). We took 1ml of the culture, washed the media out and re-suspended the culture in 1ml of YNB without aminoacids. Then, we diluted the sample in 9ml of YNB with aminoacids but without methionine for a 1:10 final dilution. We inoculated the samples for 3hrs (enough time to get GFP expression but not enough to see the fitness costs associated with this expression). We took pictures of six randomly sampled fields of view under white and UV fluorescent light at 100X magnification. Camera settings were adjusted to increase contrast between the clusters and the background and were kept constant throughout the experiment.

The culture tubes were propagated under three selection regimes for five days before the final sample was taken and then we repeated the same procedure to quantify the number of met25 GFP individuals. Before every transfer we let the tubes settle in the bench for 3 minutes (intense selection), 7 minutes (intermediate selection) and 25 minutes (relaxed selection). Then we transferred the bottom 100*μ*l into 10ml of fresh YPD and let the cultures grow for 24hrs at 30°C and 250rpm before the next transfer. We used a similar procedure to evaluate frequency-dependent fitness but this time we started with a 10:90 mix (in volume) with either a majority of “Big” with the GFP marker or “Small” and performed competition. Each competition assay was performed in four independent replicates and at least 58 individuals counted for each replicate. We had conducted similar competition assays between these two strains (data not shown) looking at changes in size distributions as proxy of frequencies. These previous trial experiments demonstrated strong patterns (consistent with the data presented here) that could be identified with small number of replicates (*n* < 5) using the actual frequencies (lower variance within treatments).

### Drafting effects

To evaluate the effect of drafting on settling at high density we started cultures mixing 0, 10, 25, 50, 75, 90 or 100*μ*l of “Big” and a the volume necessary of “Small” to complete 100*μ*l and inoculate each mix in YPD. After 24hrs we mixed well each culture and took a 10*μ*l into 990*μ*l of water to take pictures in the microscope. Pictures were taken and analyzed as described for size measurements, except only three fields of view were captured. We then took 1.5ml of culture, let it settle in the bench for 7 minutes and discarded the upper fraction leaving only 100*μ*l left. We then re-suspended the left fraction into 990*μ*l of water and after homogenizing we took 10**μ**l for dilution and microscopy. We measured the size of at least 40 individuals of each mix before and after settling. Lastly, we calculated the median size of each mix as well as the difference between the median size before and after settling.

In addition, we measured the settling speeds of individual clusters in the presence of a high density of clusters of the same strain (“Big” or “Small”) or the other strain (“Small” or “Big”). We grew each strain in six different tubes for 24hrs. After growth, we stained 2ml of each tube dark red with safranin: we took a sample of 1ml of each tube, pellet the sample and discard the supernatant, then we re-suspended the sample in water and stained with 100*μ*l of a stock solution of safranin for 10 minutes and then washed two times. For each unstained tube we took two 90*μ*l samples (background strain) and mixed one with 10*μ*l of stained “Big” and one with 100*μ*l of stained “Small” (focal strains), therefore using a block design for the different cultures. We then introduced the 1ml mixture into a borosilicate chamber of 1mm depth and 1cm width with and let it settle for 1 minute to reduce effects related to the introduction of the liquid (Fig.3C). After that we took approximately 2 minutes of video at 10 frames per second with a small digital microscope. The number of replicates and minutes of movie was determined in trial experiments while calibrating the device (data not shown). The first minute shows mainly effects of liquid introduction and after two minutes most of the cluster density has settled bellow the camera. Using FIJI [33] with ImageJ2 [34] we split the channels of the video and kept green channel for analysis, we then subtracted the background by calculating the maximum intensity z-projection and then the difference of the projection and each frame of the video. We tracked each cluster as it settle and measured the speed of the track using TrackMate [31, 33] with plain Laplacian of Gaussian segmentation (LoG) and a linear motion tracker with a maximum gap of 5 frames to allow for clusters that get shadowed by the background strain and then appear again a few frames later. We used the same filter on spot quality for all videos minimizing the detection of other particles and image noise and maximizing the detection of clusters of the focal strain (including small ones) but adjusted the track length to get tracks of most of the clusters detected but remove false positives. Lastly we manually check for real tracks and remove false positives as well as particles moving against flow because of hydrodynamic disturbances (we were only interested in settling speeds).

To analyze the track speed data we first log transformed the track speed average to meet model assumptions (Fig. S5). We performed a mixed-effects model with the log transformed average track speed as the response variable, focal strain and background as the fixed model effects and replicate as a random effect. Then we performed a set of planned contrasts to compare the difference between the settling speeds of each strain in the different backgrounds to test for the importance of drafting [29, 35]. In addition, to test for the benefit of “Big-late” in a majority “Small” background we also contrasted the speeds of both strains in a “Small” background. For these tests we used the “nlme” [36], “lme4” [37] and “multcomp” [38] packages in *R* [28].

## Aknowledgments

We would like to thank Mariah Crab for laboratory assistance and Henry Wyneken for statistical advice. Ruth Shaw, Jonathan Losos, Joan Strassmann, William Driscoll, Melanie Bowman and BIG-WIG members (especially Daniel Stanton) all provided useful feedback and thoughtful comments. This work was developed as part of MRG doctoral dissertation with support from ICGC as well as the IDF and DDF fellowships from the graduate school at the University of Minnesota. We thank R. Ford Denison and Will Harcombe for use of their microscope and plate reader, respectively. MT is funded by the John Templeton Foundation. This work was supported by NSF grant DEB-0918897. The authors have no conflicting interests to declare.

## Supplementary material

### Supplementary text

At week eight, we could only identify two groups (Big and Small, Fig.1A) when comparing the overall distributions of size. However, in previous work we had identified at least four size classes when comparing only adult sizes (excluding the small propagules) [6]. In addition each of these strains displays a slightly different settling profile (Sup). Using growth and settling rates from a strain of each of these four size groups (including our Big and Small strains) in our computer simulations, we observed that trade-offs and fitness differences between strains might also be playing a role in maintaining diversity in this system. Small fitness differences between the two small strains results in very slow displacement (more so in the absence of settling selection) with coexistence for more than the total duration of the experiment (Sup figure). More surprisingly, adding settling selection seems to be sufficient to allow for stable coexistence (in our simulations at least) between one of the Big and one of the Small strains. Non-linearity in growth and settling generates complex ecological dynamics and allow for long-term coexistence. Without settling, fast growers have an advantage and rapidly outcompete settlers. At the other extreme, intense settling selection (*i.e.* short settling times) favors the fastest settlers. Coexistence of strains displaying different strategies is possible at intermediate selection (Mean=7min, 95% CI [6.5 min, 7.5 min], Fig S2B).

### Supplementary tables

**Table S1:**
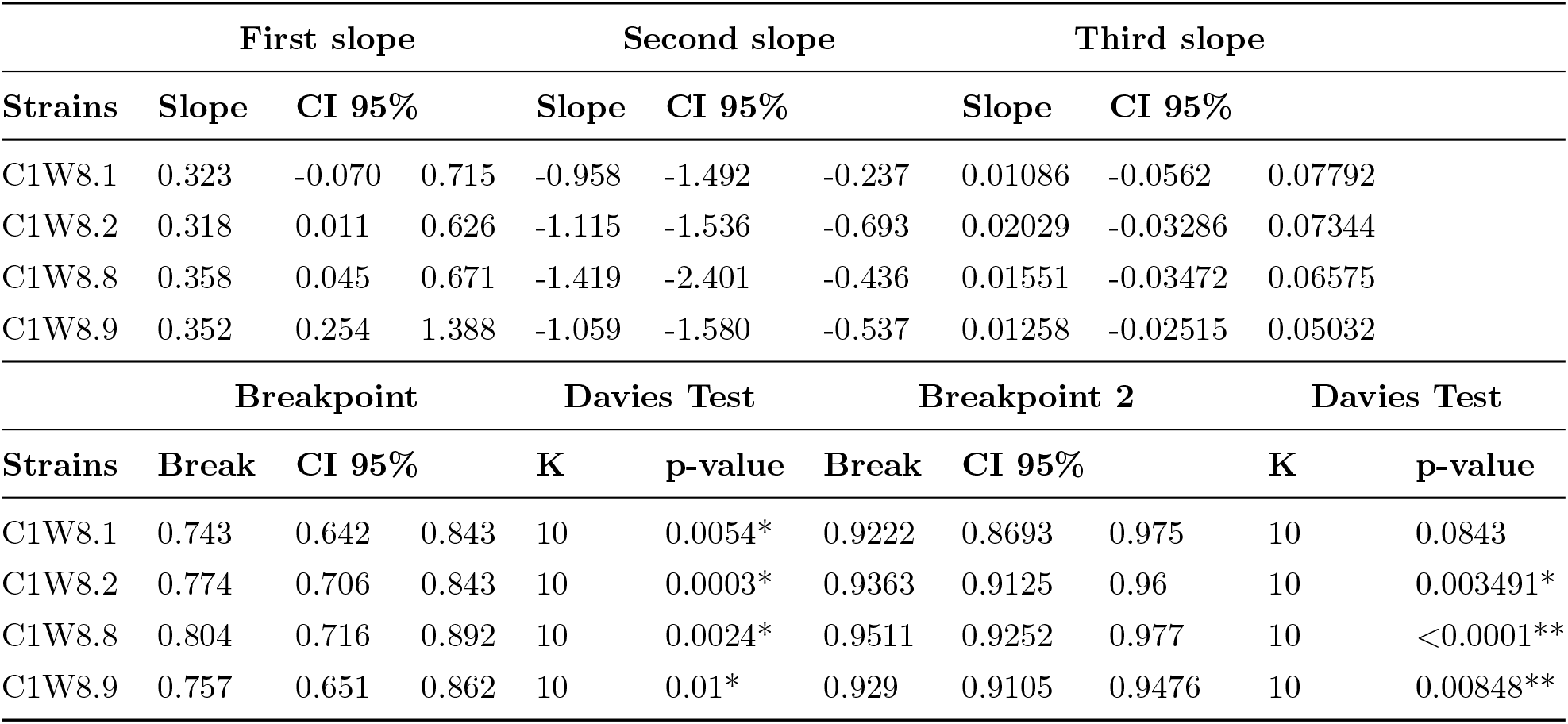
Piecewise regression of growth rate (change in optical density at 600nm over time) against optical density.

| First slope |  |  |  | Second slope |  |  | Third slope |  |  |  |
| --- | --- | --- | --- | --- | --- | --- | --- | --- | --- | --- |
| Strains | Slope | CI 95% |  | Slope | CI 95% |  | Slope | CI 95% |  |  |
| C1W8.1 | 0.323 | -0.070 | 0.715 | -0.958 | -1.492 | -0.237 | 0.01086 | -0.0562 | 0.07792 |  |
| C1W8.2 | 0.318 | 0.011 | 0.626 | -1.115 | -1.536 | -0.693 | 0.02029 | -0.03286 | 0.07344 |  |
| C1W8.8 | 0.358 | 0.045 | 0.671 | -1.419 | -2.401 | -0.436 | 0.01551 | -0.03472 | 0.06575 |  |
| C1W8.9 | 0.352 | 0.254 | 1.388 | -1.059 | -1.580 | -0.537 | 0.01258 | -0.02515 | 0.05032 |  |
| Breakpoint |  |  |  | Davies Test |  | Breakpoint 2 |  | Davies Test |  |  |
| Strains | Break | CI 95% |  | K | p-value | Break | CI 95% |  | K | p-value |
| C1W8.1 | 0.743 | 0.642 | 0.843 | 10 | 0.0054* | 0.9222 | 0.8693 | 0.975 | 10 | 0.0843 |
| C1W8.2 | 0.774 | 0.706 | 0.843 | 10 | 0.0003* | 0.9363 | 0.9125 | 0.96 | 10 | 0.003491* |
| C1W8.8 | 0.804 | 0.716 | 0.892 | 10 | 0.0024* | 0.9511 | 0.9252 | 0.977 | 10 | <0.0001** |
| C1W8.9 | 0.757 | 0.651 | 0.862 | 10 | 0.01* | 0.929 | 0.9105 | 0.9476 | 10 | 0.00848** |

**Table S2:**
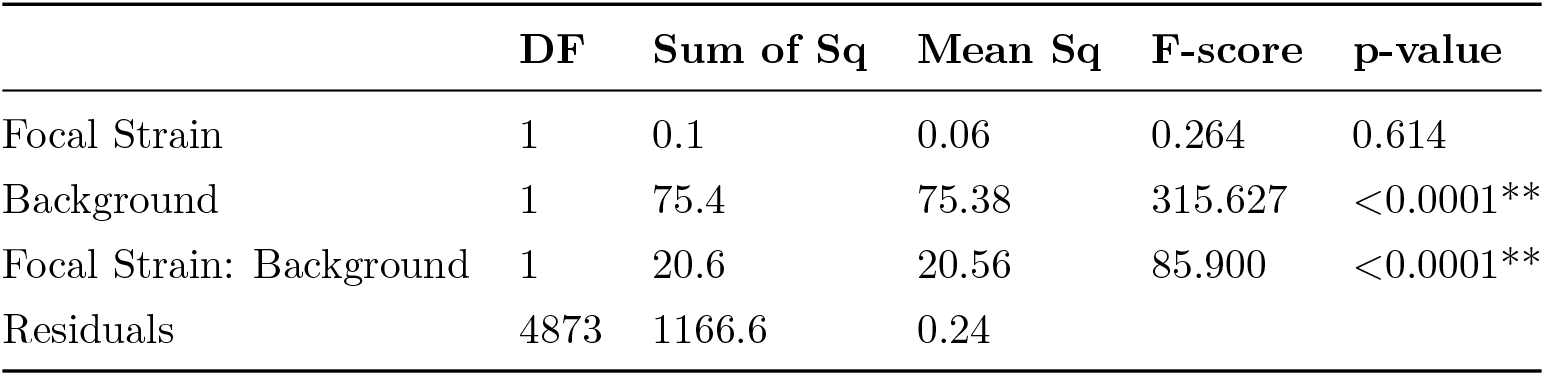
ANOVA table for mixed model effects with log transformed track settling speed as the response variable, focal strain and background (see figure 4C) as fixed effects and replicate as a random effect.

|  | DF | Sum of Sq | Mean Sq | F-score | p-value |
| --- | --- | --- | --- | --- | --- |
| Focal Strain | 1 | 0.1 | 0.06 | 0.264 | 0.614 |
| Background | 1 | 75.4 | 75.38 | 315.627 | <0.0001** |
| Focal Strain: Background | 1 | 20.6 | 20.56 | 85.900 | <0.0001** |
| Residuals | 4873 | 1166.6 | 0.24 |  |  |

### Other supplementary materials

**Video S1.** Time-lapse of the “Big-late” (C1W8-2) and “Small-early” (C1W8-9) genotypes growing from a small propagules over ten hours and 21 minutes with pictures taken every three minutes. The “Small-early”genotype starts dividing almost immediately whereas the “Big-late” genotype first grows and as cell divisions slow down it starts dividing.

**Software S1.** R-code to analyze cluster sizes in order to obtain the distributions of weighted volume or biomass proportion.

**Software S2.** R-code of competition simulations.

### Supplementary figures

**Figure S1:**
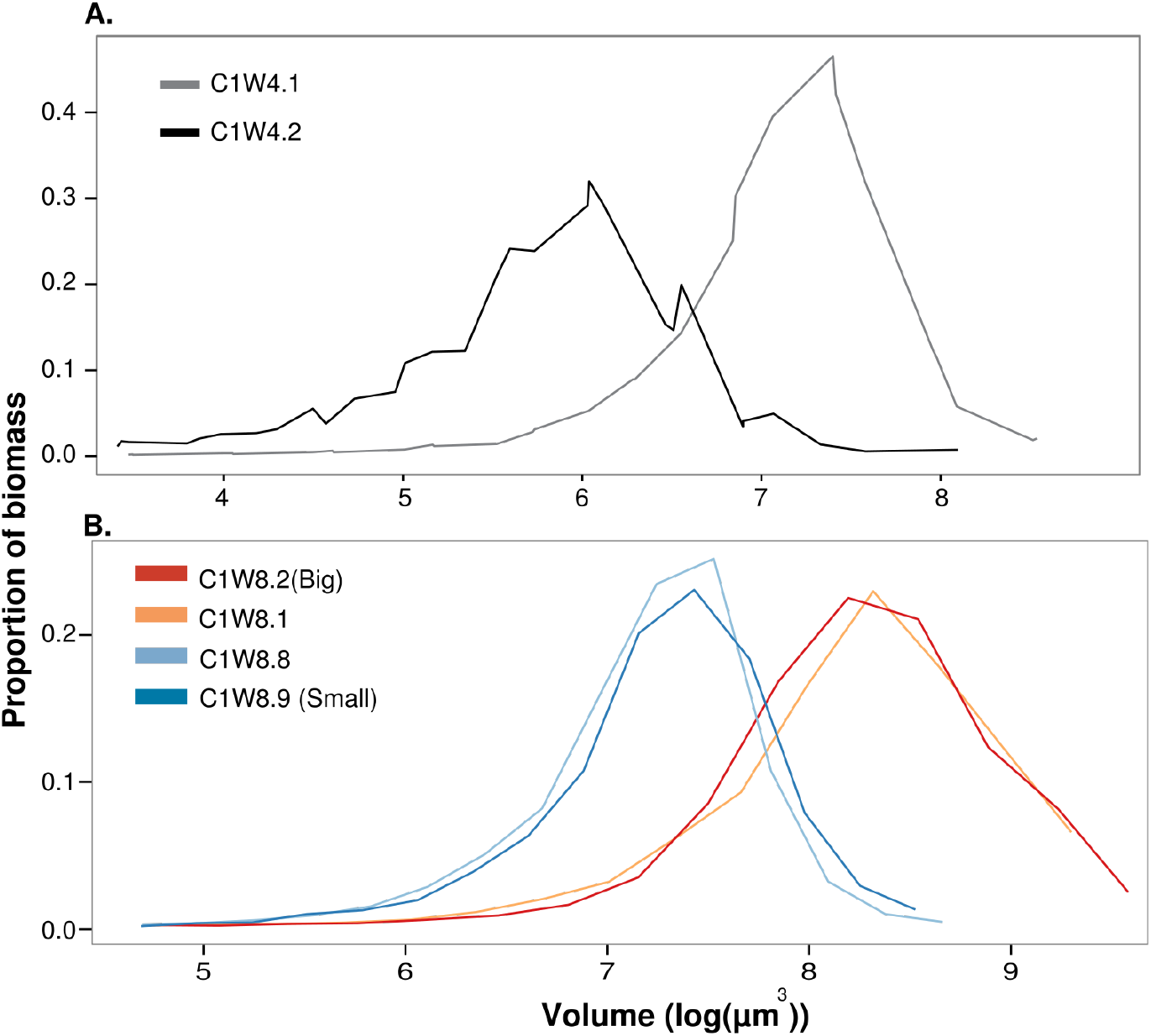
Biomass distribution of representative isolates after four (A) and eight weeks of settling selection (B). We identified size divergence in multicellular isolates, in one of the replicate populations (C1), after four and eight weeks of evolution under settling selection. At week 4, small isolates have a fairly symmetric distribution of sizes (C1W4.1), where mid-size clusters comprise most of the total biomass. Bigger isolates, instead, evolved a few larger adults taking up a large proportion of all the biomass and many very small propagules with a minimal contribution to the overall biomass (C1W4.2). Similarly large adults with small propagules are present still after 8 weeks (Small, C1W8.8), but at this time new isolates with increasingly asymmetrical distributions had evolved (Big, C1W8.1).

**Figure S2:**
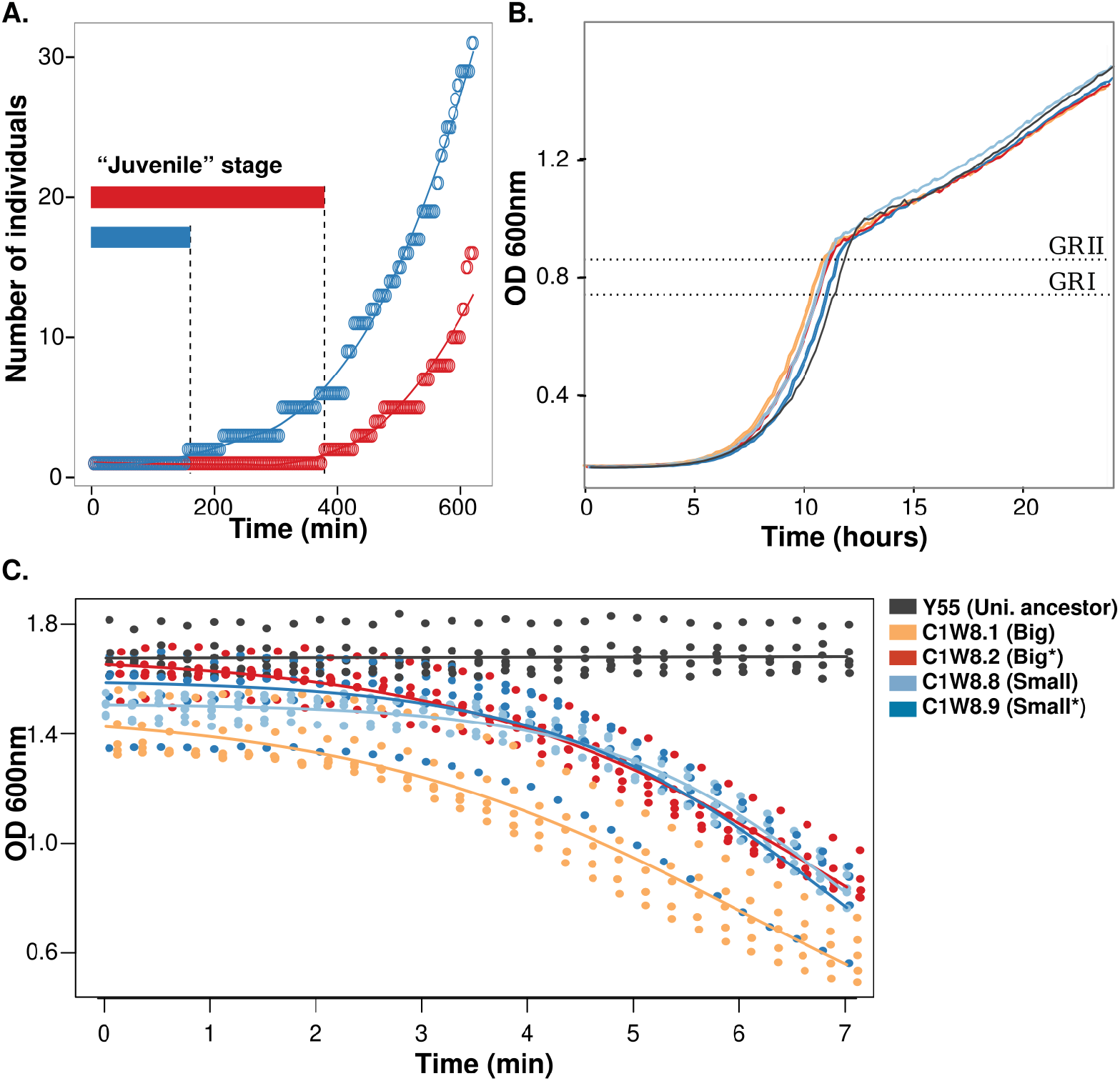
Differences in growth and settling between different strains after 60 days of selection. (A) Growth curves starting with a single multicellular individual of similar size (a small propagule). Strains with larger adult sizes have small propagules and take a long time to reach a “reproductive size”. At that point they start breaking into smaller propagules. Therefore, these propagules have a larger juvenile phase (length of time before their first reproductive event) than grower strains with smaller adults. Observations used in making this graph are recorded in Video S1. (B) Growth curves over 24hrs for the different multicellular isolates of week eight and their unicellular ancestor (gray line) as a control for growth dynamics. Dotted lines mark the minimal optical density at which at least one of the strains displays a changed in growth rate. These values were used in the simulations to model changes in growth rate during competition (Fig.3). (C) Settling dynamics of different isolates at week eight as well as their unicellular ancestor for comparison. The graph shows the reduction in optical density at 600nm as individuals settle over seven minutes. Points are data from six replicate assays. The dark gray line shows the linear fit of settling data from unicellular ancestor; the other lines represent the fitted logistic models to our data (see main text).

**Figure S3:**
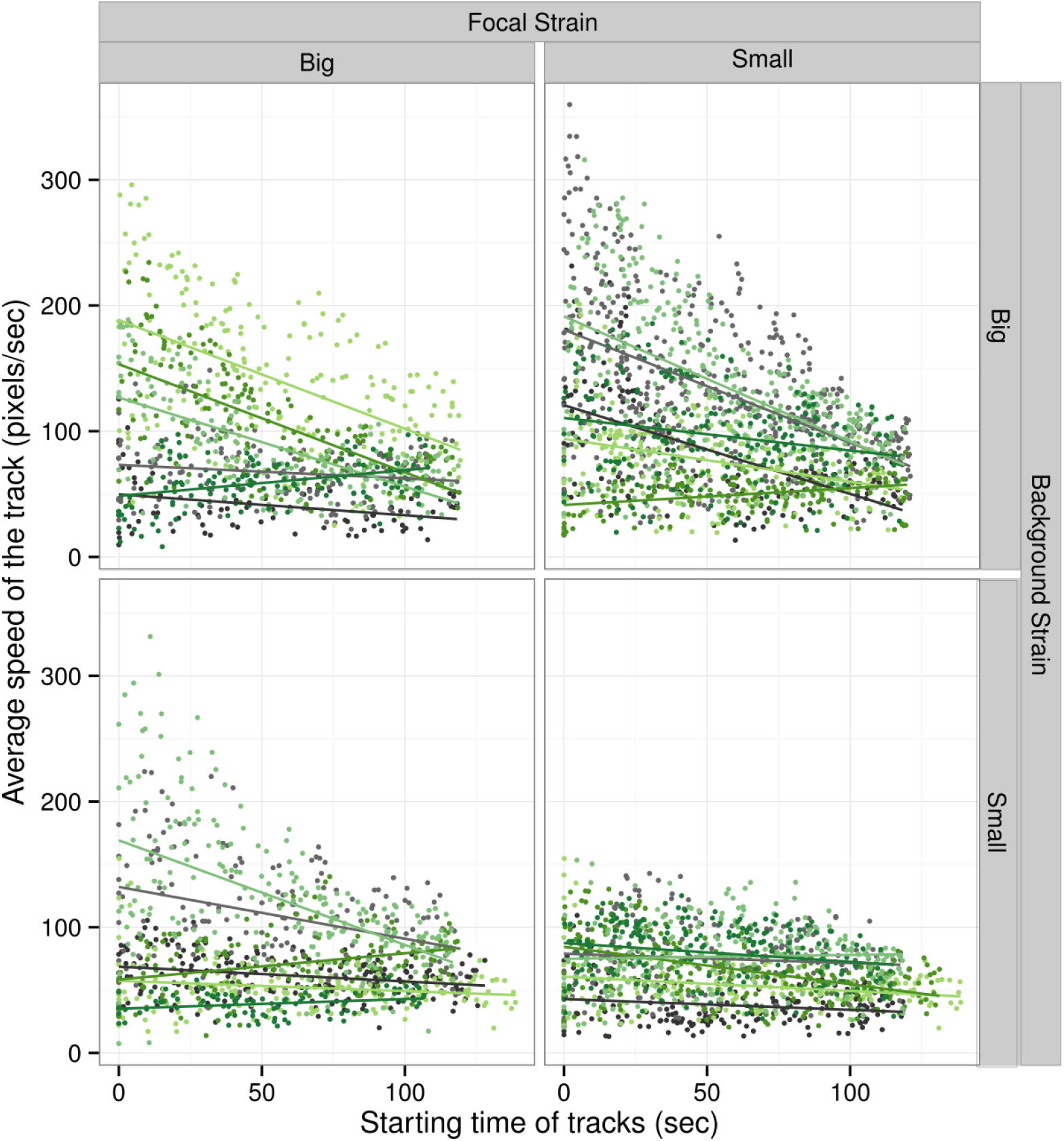
Average settling speed of clusters over recorded time ( 2min). There is more variation on the big background within and between replicates (different shades of grays and greens) as expected from their wide size distribution (where there is a small number of really big clusters that settle fast and are capable of dragging smaller clusters) and a large number of small propagules with slow settling rates.

**Figure S4:**
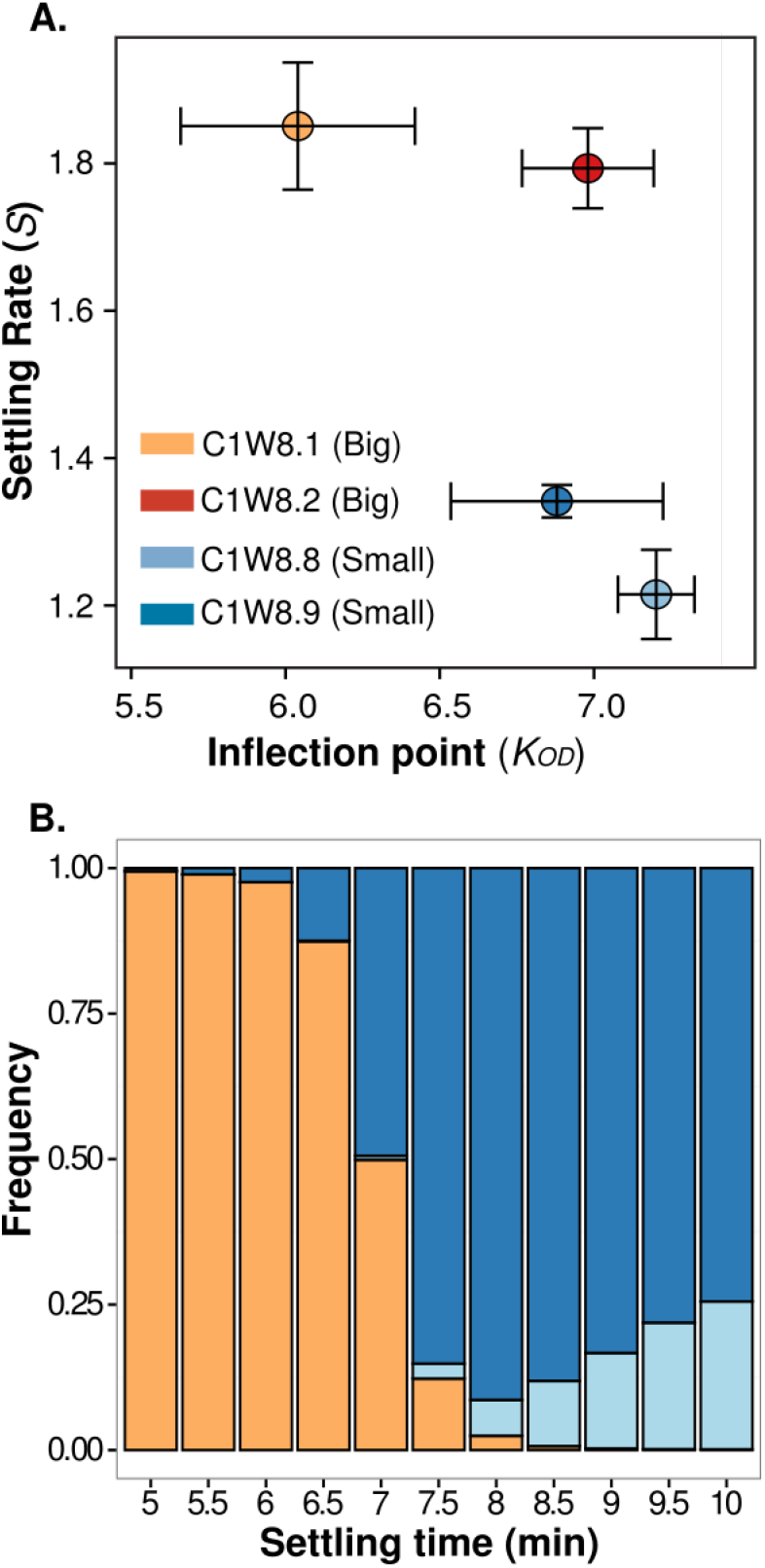
Trade-off between growth and settling allows coexistence of different strategies in simulations. (A) Settling parameters of different strains. The error bars show the standard deviation across six replicates. (B) The results of a round of simulations over 100 days of competition with 7 minutes as the settling selection favoring coexistence. Differences in breakpoint estimates resulted in differences on the selection strength (settling time) necessary for coexistence between the fastest growing and fastest settling strains, we repeated the simulations 100 times (re-calculating breakpoints in each case). In each case we were able to found a point of coexistence (Mean=7min, 95% CI [6.5 min, 7.5 min]

**Figure S5:**
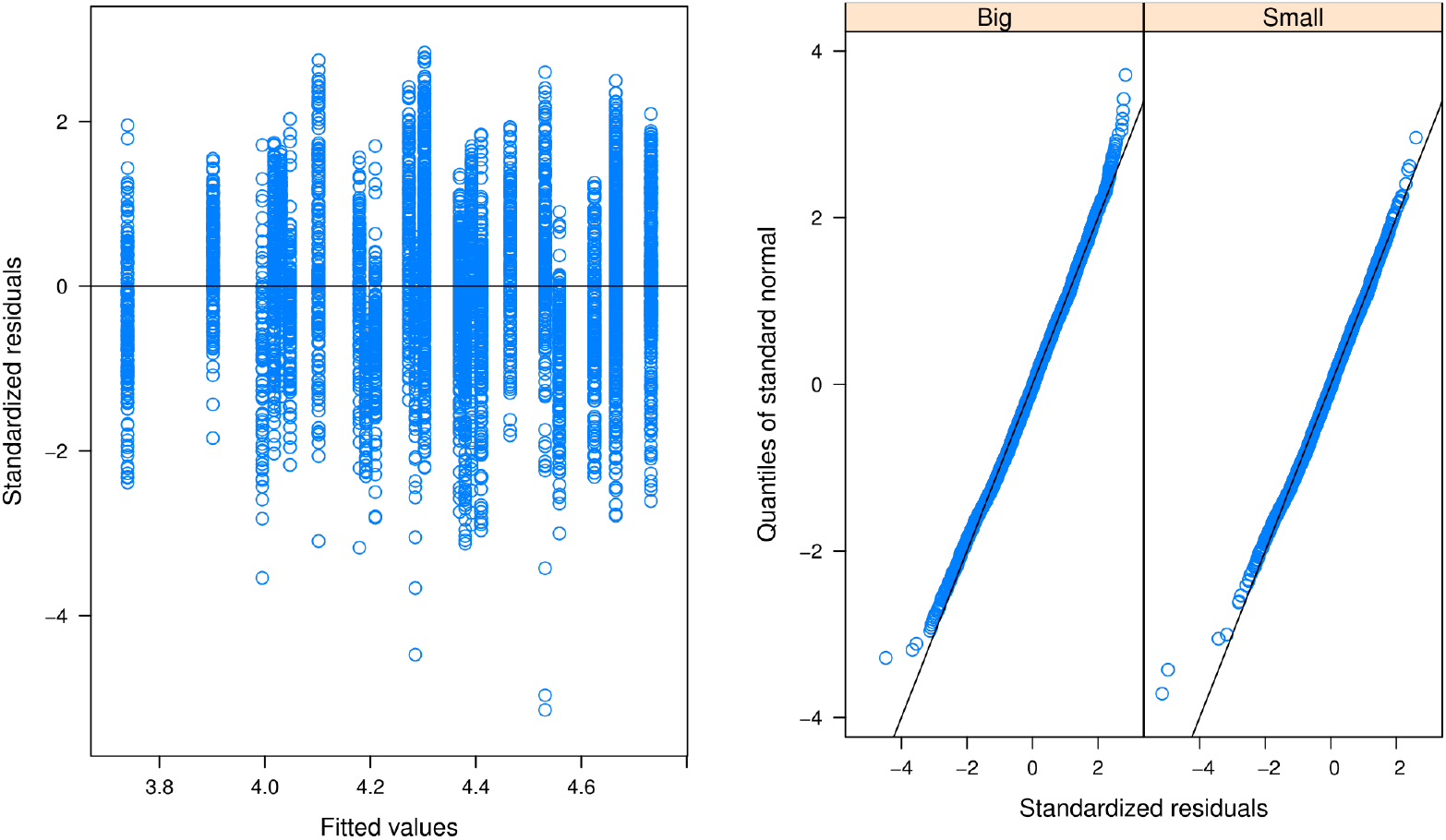
After log transformation data meets the assumption of homogeneity of variance and only deviates from normality at the extremes.

